# DiCoLo: Integration-free and cluster-free detection of localized differential gene co-expression in single-cell data

**DOI:** 10.1101/2025.11.23.689932

**Authors:** Ruiqi Li, Junchen Yang, Pei-Chun Su, Ariel Jaffe, Ofir Lindenbaum, Yuval Kluger

## Abstract

Detecting changes in gene coordination patterns between biological conditions and identifying the cell populations in which these changes occur are key challenges in single-cell analysis. Existing approaches often compare gene co-expression between predefined cell clusters or rely on aligning cells across conditions. These strategies can be suboptimal when changes occur within small subpopulations or when batch effects obscure the underlying biological signal. To address these challenges, we introduce DiCoLo, a framework that identifies genes exhibiting differential co-localization, defined as changes in coordinated expression within localized cell neighborhoods–subsets of highly similar cells in the transcriptomic space. Importantly, DiCoLo does not rely on cell clustering or cross-condition alignment. For each condition, DiCoLo constructs a gene graph using Optimal Transport distances that reflect gene co-localization patterns across the cell manifold. Then, it identifies differential gene programs by detecting changes in connectivity patterns between the gene graphs. We show that DiCoLo robustly identifies differential gene co-localization even under weak signals or complex batch effects, outperforming existing methods across multiple benchmark datasets. When applied to mouse hair follicle development data, DiCoLo reveals coordinated gene programs and emerging cell populations driven by perturbations in morphogen signaling that underlie dermal condensate differentiation. Overall, these results establish DiCoLo as a powerful framework for uncovering localized differential transcriptional coordinated patterns in single-cell data.

## 1 Introduction

Single-cell RNA sequencing (scRNA-seq) enables transcriptome-wide profiling of thousands of individual cells, revealing heterogeneity among cell populations, including distinct cell types and subtypes, continuous transitions, and rare populations [1]. A central challenge in scRNA-seq analysis is to uncover how genes coordinate their expression within specific cellular contexts. Detecting coordination patterns is essential for understanding regulatory programs defining cellular states and functions [2–5]. Of particular interest are cases where gene coordination changes across biological or disease conditions. Such changes can arise from pathway activation or silencing within specific cell subpopulations [5], the disruption of regulatory programs under perturbation [6], or the emergence and loss of specialized cell populations [7, 8]. Identifying differential gene coordination patterns within cell subpopulations provides insight into how fine-grained transcriptional regulation shifts across conditions.

Despite its importance, detecting changes in coordination patterns remains a statistical and computational challenge. The difficulty arises from the large number of gene pairs and the substantial technical noise and sparsity that characterize scRNA-seq [9]. A common approach to addressing this task is differential co-expression analysis, where a measure of statistical dependence between gene pairs is computed separately for each condition or cell cluster. The correlation matrices are then compared in order to detect gene pairs or modules with altered correlations [2]. Among commonly used tools, DiffCoEx [10] compares correlation matrices between conditions to cluster genes that exhibit differential correlation. DGCA [11] applies permutation testing to identify gene pairs with significant differential correlations. However, the large number of pairwise tests significantly reduces statistical power. In addition, these methods often rely on correlation measures (e.g., linear or Spearman correlation) within predefined cell clusters. As a result, they may miss localized changes that occur, for example, within a small cell subpopulation or at specific cell states within a continuous trajectory.

Cluster-free differential analysis methods, such as miloDE [12] and LEMUR [13], identify gene-level differences between conditions without relying on predefined cell clustering. These methods first integrate cells from all conditions into a common low-dimensional space and test for differential expression within cell neighborhoods. Thus, they enable the discovery of transcriptional changes within local cellular structures. The main limitation of this approach is the need to integrate cells from different conditions. This step may be challenging when strong batch effects or significant transcriptomic shifts cause the cell manifolds to separate substantially. Applying batch-correction procedures may distort the true biological signals and obscure condition-specific cellular structures [14, 15].

A gap remains in identifying changes in gene coordination patterns between conditions, that occur within specific regions of the cell manifold. To address this gap, we propose DiCoLo (**Di**fferential **Co**-**Lo**calization analysis), a framework that captures genes exhibiting differential coordinated expression between conditions within localized cell neighborhoods, without requiring predefined cell clusters or cross-condition cell integration. DiCoLo quantifies gene–gene *co-localization*, the coordination between genes within local neighborhoods. Co-localization is computed in each condition using the Optimal Transport (OT) distance [16], which measures the minimal cost of transforming the distribution of one gene’s expression into another across the cell manifold. OT penalizes the moving of mass between distant cells more than between neighboring cells. Thus, pairs of genes activated within overlapping cellular neighborhoods have small OT distances, whereas genes expressed in distal subpopulations of cells yield large distances [8]. This property makes OT sensitive to co-localization patterns confined to specific regions of the manifold. Based on the OT distance measure, DiCoLo computes a gene graph for each condition, where edge weights are set according to the co-localization between genes. Detecting differential gene programs is framed as comparing two graphs to identify groups of vertices (genes) with different connectivity profiles. This is achieved through a differential graph operator that contrasts the spectra of the two graphs. By computing gene graphs separately for each condition, DiCoLo bypasses the risk of signal distortion and over-correction common to integration-based methods.

To evaluate our approach, we first apply DiCoLo to a human PBMC dataset with simulated condition-specific co-localized gene groups, demonstrating that our method recovers the known differential co-localization signals. We further benchmark DiCoLo against differential co-expression and cluster-free differential expression methods across diverse simulation settings, demonstrating that DiCoLo effectively detects differential co-localization even under weak signals and remains robust in the presence of strong batch effects. Finally, we apply DiCoLo to mouse embryonic skin datasets with distinct morphogen mutants, identifying condition-specific co-localized gene modules that reveal transcriptional programs and cell populations driven by morphogen signaling. Together, these results highlight the utility of DiCoLo as a cluster-free, integration-free framework for detecting differential co-localization and its biological significance.

## 2 Results

### 2.1 Overview of DiCoLo

DiCoLo identifies genes whose co-localization patterns differ between conditions through a three-step workflow, illustrated in Fig. 1. The three steps are explained in detail in Section 4.

**Fig. 1.**
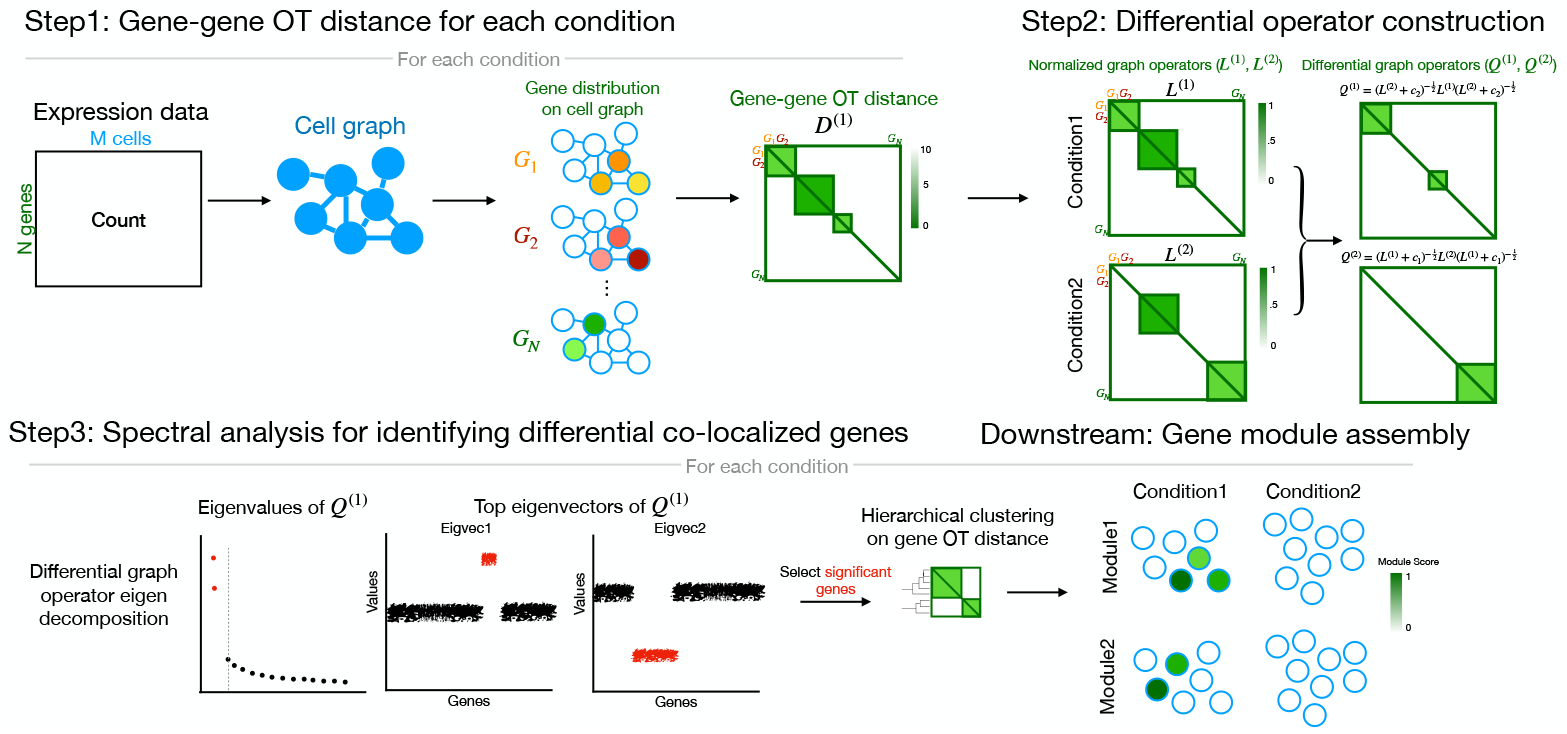
Schematic overview of the DiCoLo workflow. Step 1: For each condition, gene-gene OT distances are computed based on a cost function determined by a cell graph. Step 2: OT distances are converted into normalized graph operators, which are contrasted in two directions to construct differential operators for detecting co-localized genes unique to condition 1 (*Q*^(1)^) and condition 2 (*Q*^(2)^). Step 3: Spectral decomposition of the differential operator identifies genes with differential co-localization patterns. Downstream: Differentially co-localized genes are grouped into modules by hierarchical clustering, and the module scores (mean expression across module gene) are mapped onto cell embeddings from each condition.

#### Step 1: Computing the OT distance between the expression level of pairs of genes

DiCoLo constructs a gene-gene distance matrix for each condition. This process begins by constructing a cell graph for each condition from the transcriptomic profile of the cells. The graph defines a *cost* function that reflects the distances between cells along the manifold. Drawing upon the graph-based OT distance formulation [8], each gene’s expression profile is normalized into a probabilistic distribution over the cell graph, and pairwise OT distances are computed between these gene distributions to form a gene-gene distance matrix for each condition.

#### Step 2: Constructing differential graph operators

For each condition, DiCoLo constructs a graph whose nodes correspond to the genes and whose weights are functions of the OT distances computed in Step 1. We then search for sets of nodes (genes) that are densely connected in one graph but not in the other. Inspired by cross-graph comparison methods [17–20], DiCoLo computes two directional differential graph operators: each capturing the node connectivity structures that are distinctly strong in one condition relative to the other.

#### Step 3: Spectral analysis for identifying differential co-localized genes

From the differential graph operators, DiCoLo identifies sets of genes whose co-localization structure differs between conditions through eigen-decomposition. Significant eigenvectors are selected using the knee point of the eigenvalue spectrum, following the commonly used practice for selecting informative components[21, 22]. Genes with large absolute loadings in these eigenvectors correspond to those showing differential co-localization across conditions[18]. To reveal broader coordinated gene programs, these genes can be grouped into co-localized modules based on their OT distances from Step 1. Aggregating gene expression within each module allows us to map transcriptional programs to specific cell populations, while the module composition facilitates downstream pathway analysis for biological interpretation.

### 2.2 Demonstration of DiCoLo on a human PBMC dataset

For illustration, we apply DiCoLo in a controlled scenario using a real PBMC dataset with simulated co-localization signals. We randomly divided CD14^+^ monocytes into two biologically identical subsets, Sample 1 and Sample 2 (Fig. 2A). We generate six synthetic co-localized gene modules: (i) *Sample-specific modules:* Two gene modules, termed M-S1a and M-S1b, whose expression is confined to distinct neighborhoods N-S1a and N-S1b in sample1. Similarly, modules M-S2a and M-S2b appear only in sample 2 in cell neighborhoods N-S2a and N-S2b. (ii) Two common gene modules, M-Ca and M-Cb, co-localized in shared neighborhoods N-Ca and N-Cb containing cells from both samples. All other genes retain their original expression in both samples. Details of the simulation procedure are provided in the Methods section. Our goal is to correctly identify sample-specific modules (e.g., M-S1a) while filtering out common modules (e.g., M-Ca).

**Fig. 2.**
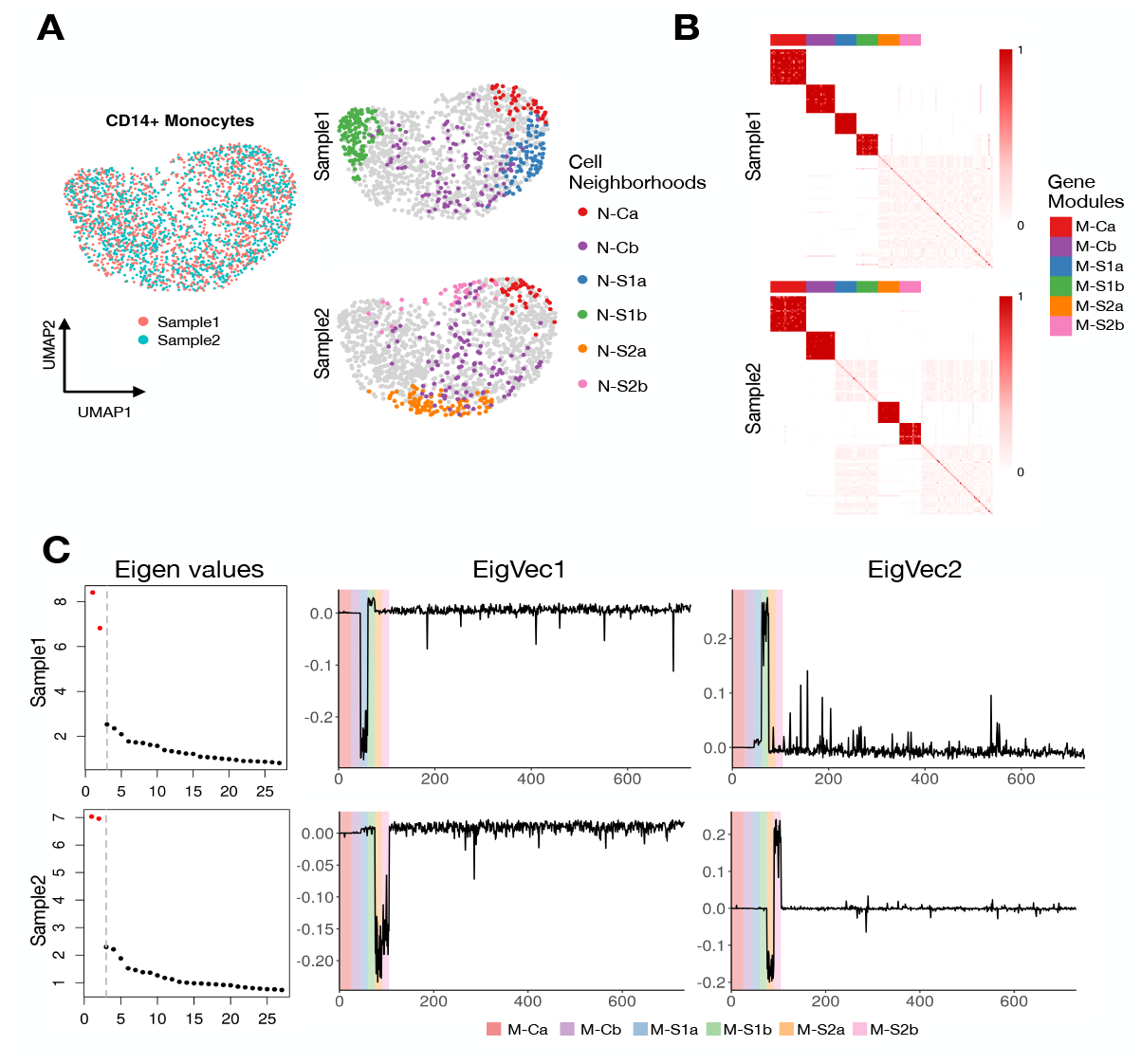
Illustration of DiCoLo using human pbmc10k dataset with artificial co-localized signals. (A) Left: UMAP of all cells, colored by sample. Right: The same UMAPs, split by sample, showing four sample-specific (N-S1a, N-S1b in Sample1; N-S2a, N-S2b in Sample2) and two common cell neighborhoods (N-Ca, N-Cb). The latter contains cells from both samples. (B) Gene-gene normalized graph operators for Sample 1 (top) and Sample 2 (bottom), min-max scaled for visualization. Genes are ordered by module membership: the six simulated modules are shown first, followed by 50 randomly selected genes. Color bars indicate module identity. (C) Eigen-decomposition of the differential operators. Left: Eigenvalue spectra for Sample 1 (top) and Sample 2 (bottom); significant eigenvalues are highlighted in red. Right: Eigenvector loadings for the top two eigenvectors. Genes are ordered by module membership, with the background colors indicating module identity.

Fig. 2B visualizes the normalized gene-graph operator computed in Step 2. Sample-specific modules form dense diagonal blocks in their respective samples: M-S1a and M-S1b appear only in Sample 1, while M-S2a and M-S2b appear only in Sample 2. In contrast, common modules (M-Ca and M-Cb) form dense co-localized blocks in both samples. Fig. 2C shows the eigenvectors and eigenvalues of the two differential graph operators computed in Step 3. The first two eigenvalues of both operators are substantially larger than the remaining eigenvalues, indicating that each sample contains two major differential signals. The two corresponding eigenvectors have large absolute loadings on genes in the sample-specific modules. In contrast, the common modules and the rest of the genes show negligible loadings. This demonstrates that DiCoLo correctly identifies condition-specific co-localized gene modules while ignoring shared patterns.

### 2.3 Benchmarking DiCoLo across diverse single-cell scenarios

To systematically evaluate DiCoLo’s performance, we designed simulations across three real scRNA-seq datasets that differ in cell type, sequencing platform, read depth, and the presence of batch effects (Fig.S1, Supplementary Data 1, Methods - ‘Dataset and Processing’). For each dataset, we simulate one co-localized gene module in one condition while leaving the other unchanged (Methods – ‘Simulation workflow’). We then test whether DiCoLo and competing methods can correctly identify these genes while avoiding false discoveries driven by batch effects or biological variation. We leverage existing batch effects in real data rather than simulating artificial technical variation [15], allowing us to test robustness under realistic conditions. The first dataset (human pbmc10k, Fig. S1A) provides a baseline with no batch effect, using the CD14^+^ monocytes described above, randomly split into two subsets. The second dataset (human pbmcsca, Fig.S1B) introduces inter-donor biological variation, using alpha cells from two healthy donors of a human pancreas dataset [23] (human2 and human3), both generated using the inDrops platform by the same laboratory. The third dataset introduces cross-platform technical variation (human pancreas, Fig. S1C), using cytotoxic T cells from the same donor sequenced with two different technologies [24] (Drop-seq and inDrops), which differ substantially in sequencing depth (median nUMI = 1207 vs. 272; Supplementary Data 1).

We compare DiCoLo against existing methods for detecting differential patterns: DGCA[11] for differential co-expression analysis, and miloDE[12] and LEMUR[13] for cluster-free differential expression. As no direct methods currently exist for differential co-localization, we utilize miloDE and LEMUR as the closest available baselines for detecting localized condition-specific signals without predefined clusters. Each method ranks genes by the significance of their differential signal (see Methods - ‘Benchmarking with existing methods’). We evaluate performance using normalized area under the precision–recall curve (normalized AUPRC), which accounts for class imbalance [25] (see Methods). We test three key parameters that affect signal detectability: (1) neighborhood size, which controls the spatial extent of localized gene activation (Fig. 3A); (2) dropout rate, which controls signal sparsity (Fig. 3B); and (3) the number of perturbed genes (Fig. 3C). When varying one parameter, the others are held constant (neighborhood size = 10% of the total cells, dropout rate = 0.4, number of genes = 15).

**Fig. 3.**
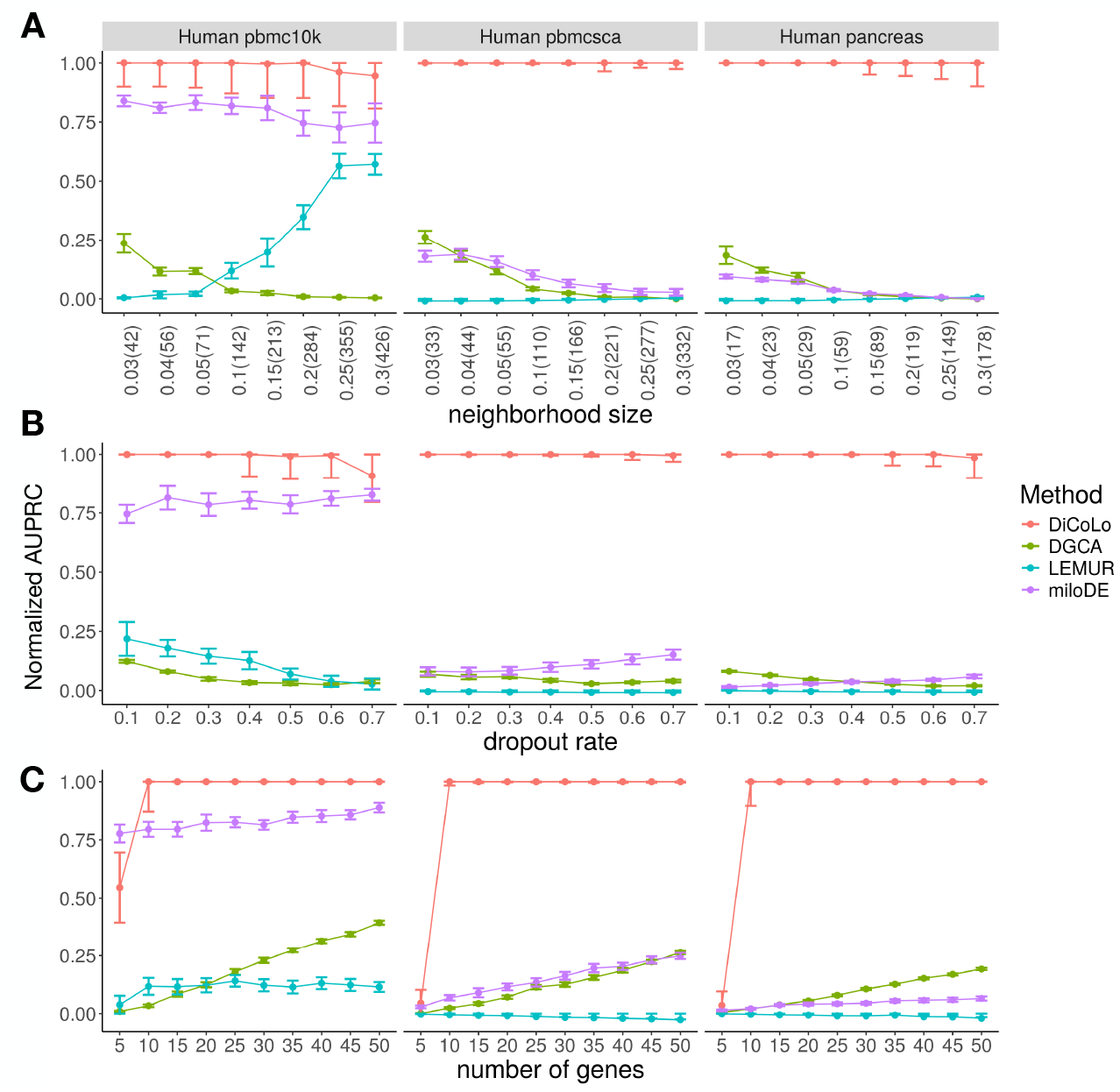
Benchmarking DiCoLo across simulated datasets. (A-C) Performance comparison (normalized AUPRC) of DiCoLo with DGCA, LEMUR, and miloDE under three simulation parameters across three datasets (human pbmc10k, pbmcsca, and pancreas). Each point represents the mean ± s.e.m. over 20 simulations (signal simulated in each condition 10 times). (A) neighborhood size, shown as percentage with absolute cell number (3–30% of the total cells), (B) dropout rate (0.1–0.7), (C) number of perturbed genes (5–50).

DiCoLo consistently outperforms all competing methods across simulation settings (Fig. 3). It remains robust performance across a wide range of neighborhood sizes, dropout rates, and number of co-localized genes. The only exception occurs with extremely weak signals (number of perturbed genes = 5), where all methods perform comparably (Fig. 3C). miloDE performs well when batch effects are minimal (human pbmc10k dataset) but shows reduced accuracy under stronger batch variation. DGCA and LEMUR show consistently lower performance across all settings. These results demonstrate that DiCoLo provides more reliable detection of differential co-localization signals even under challenging conditions with weak signals and strong batch effects.

### 2.4 DiCoLo identify condition-specific co-localized patterns in mouse dermal condensate genesis

Mouse hair follicle dermal condensates (DCs) arise within the dermis around embryonic day 14.5 (E14.5) and play an essential role in initiating hair follicle morphogenesis[7]. DC differentiation is regulated by the coordinated activity of the Wnt/*β*-catenin and Sonic Hedgehog (SHH) signaling pathways[7, 26, 27]. To investigate how these pathways drive DC differentiation, we apply DiCoLo to compare genetic mutants with altered Wnt or SHH signaling against wildtype controls. We aim to identify gene modules exhibiting differential co-localization within specific cell populations, thereby revealing regulatory programs that emerge, shift, or disappear in response to pathway perturbation.

#### Premature SHH activation drives precocious DC and quiescence programs

We first examine the SmoM2 mutant (Fig. 4A), which induces high SHH activation across dermal cells at E13.5, a stage preceding natural DC formation[7]. At this stage, wildtype controls (CTL) exhibit a Wnt signaling gradient in the upper dermis (UD) but have not yet initiated SHH expression or DC formation. DiCoLo identifies three gene modules that are co-localized in the SmoM2 mutant but dispersed in the CTL (Fig. 4, S2, all genes in Supplementary Data 2). SmoM2-M1 contains canonical DC markers such as *Sox2* and *Gal*. This module captures the SHH-driven, precocious emergence of DC populations that are absent in the CTL at this stage. SmoM2-M2 comprises genes enriched in the p53 pathway (Fig. 4C), including *Trp53inp1*, which is involved in cell cycle exit. The co-localization of these genes defines a SmoM2-specific quiescent cell population, which is a critical intermediate cell state required for terminal DC differentiation[28, 29]. This finding unmasks the role of SHH signaling in regulating cell cycle exit during DC development. SmoM2-M3 contains Wnt signaling genes (e.g., *Robo2, Bmp4*, and *Wif1*) and DC markers (e.g., *Foxd1, Dclk1*, and *Sox18*) that are co-localized across a broader region of the UD in the SmoM2 mutant. This module indicates that high SHH activation within the Wnt-active UD region induces DC gene programs in a broader range of cells, suggesting a premature interaction between these two pathways in the mutant.

**Fig. 4.**
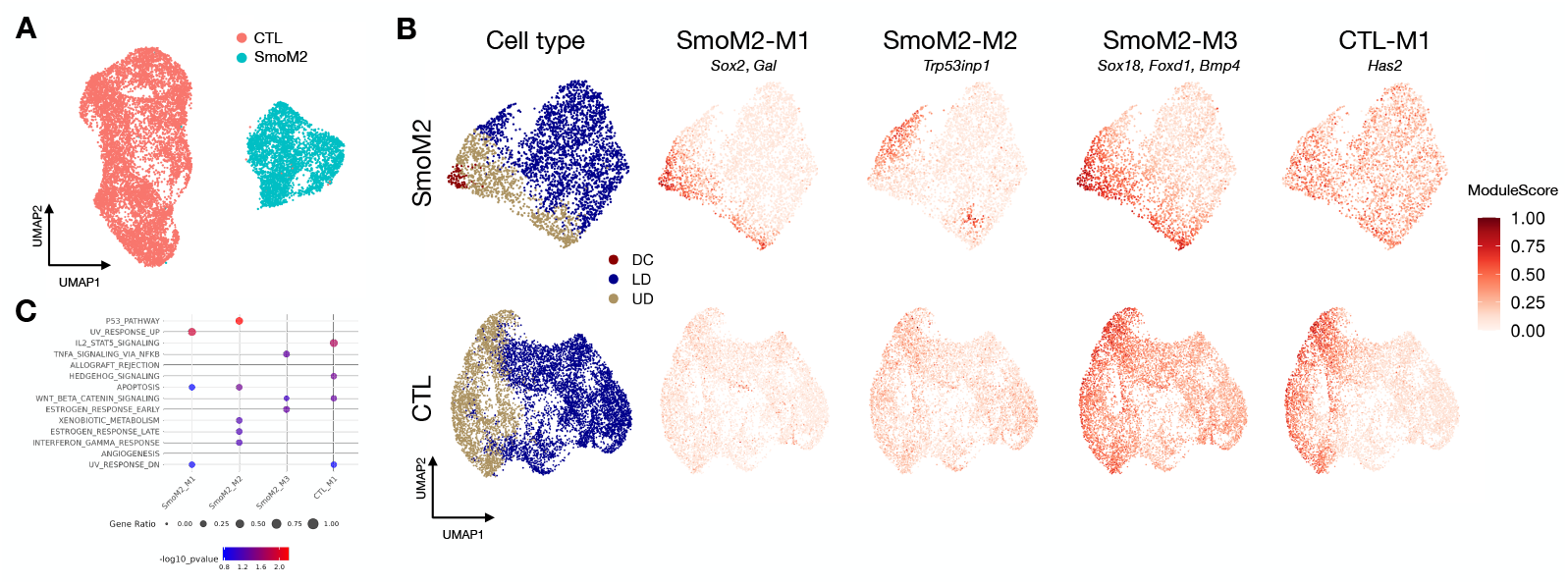
DiCoLo reveals condition-specific co-localized gene modules in DC genesis (SmoM2 vs wild-type control). UMAP of cells color-coded by condition. (B) UMAP of cells embedded separately for each condition, color-coded by cell type (left) and by co-localized gene modules (right), including three SmoM2-specific and one control-specific modules. (C) MsigDB pathway enrichment analysis of the four gene modules, showing the top 5 enriched pathways (*p* value < 0.1) for each module.

#### Widespread SHH disrupts UD-confined gene programs

Modules unique to the CTL capture gene programs confined to a specific cell region due to localized signaling, which are disrupted in the mutant background. For example, CTL-M1 contains genes co-localized in the UD of CTL embryos (Fig. 4, S3), where Wnt and other UD-related signals are active before SHH activation. This co-localization pattern is disrupted in SmoM2, where SHH signaling becomes broadly activated throughout the dermis. This loss of co-localization suggests that widespread SHH activation breaks down gene programs confined to the UD, which are likely regulated by Wnt or other UD-asscoiated signals. Notably, several genes in this module (e.g. *Has2* and *Runx3*) are direct SHH targets, indicating that SHH pathway activity may be primed by Wnt signaling at E13.5 even before SHH ligand expression begins.

#### Wnt deficiency prevents DC formation and pathway coordination

To evaluate the role of Wnt signaling, we apply DiCoLo to the Wntless (Wls) mutant, which results in deficient Wnt signaling and inhibits DC formation[8, 30]. At E14.5, when DCs are normally present, DiCoLo identifies two modules co-localized in CTL but disrupted in the Wls condition (Fig. S4, S5). CTL-M1 contains canonical DC markers (e.g. *Sox2* and *Gal*), confirming that DC populations fail to form without proper Wls signaling. CTL-M2 is enriched for both Wnt and SHH signaling pathway genes (Fig. S4C), including the Wnt target gene (*Lef1*), SHH target genes (*Ptch1, Ptch2*), and the DC-specific cell-cycle inhibitor *Cdkn1a*. These genes coordinate in the UD and DC populations of CTL embryos at E14.5[30, 31], but lose their coordinated expression pattern in the Wls mutant, demonstrating that Wnt activity before DC formation is required for subsequent SHH activation.

Together, these analyses demonstrate how DiCoLo reveals condition-specific gene coordination patterns within cell subpopulations along continuous developmental systems with strong transcriptomic shifts between conditions—patterns that would be difficult to detect through standard clustering-based co-expression analysis or integration-based differential expression methods.

## 3 Discussion

We present DiCoLo, a framework for detecting differential co-localization of genes between two conditions without requiring explicit cell alignment across conditions. This is particularly valuable when significant batch effects or substantial biological shifts make alignment unreliable. The core idea of DiCoLo is to model gene-gene relationships using optimal-transport-based graphs within each condition, and then compare these graphs using a Laplacian operator. This operator highlights subsets of genes that are co-localized in one condition but not in another, revealing a distinct mode of gene-gene variation that cannot be captured by conventional integration-based differential expression or correlation analyses.

Using simulations based on real datasets with realistic batch effects, we demonstrate that DiCoLo robustly identifies condition-specific co-localized gene groups even when signals are weak and obscured by strong batch effects. In scenarios where only a few genes are perturbed or dropout rates are high, and where large batch effects exist between two conditions, DiCoLo remains consistently accurate and outperforms existing methods, filling an analytical gap in detecting coordinated gene changes within local cellular neighborhoods.

Beyond synthetic benchmarks, we apply DiCoLo to real mouse embryonic skin datasets to demonstrate its ability to uncover biologically meaningful differential co-localization patterns. In the SmoM2 and Wls morphogen signaling mutants, DiCoLo identifies condition-specific co-localized gene modules that reveal transcriptional programs emerging or diminishing under pathway perturbations. Critically, these real biological patterns exhibit the challenging characteristics our benchmarks are designed to address: localized coordination confined to small cell subpopulations within continuous, gradient-like developmental trajectories, combined with substantial transcriptomic shifts between conditions –a combination that confounds both clustering-based co-expression and integration-based differential expression methods. These applications validate DiCoLo’s ability to capture subtle yet biologically meaningful coordination changes in practical experimental settings, bridging the gap between methodological benchmarking and biological discovery. It is important to note the biological scope imposed by the optimal transport framework. DiCoLo is explicitly designed to identify localized co-expression within subpopulations rather than global gradients. Broadly expressed genes (e.g., housekeeping genes or cell-cycle genes) often exhibit uniform distributions that yield low transport costs; therefore, we apply an expression threshold (default genes expressed in ¡50% of cells) to filter these uninformative cases. Within this localized regime, the framework proves highly robust: it detects signals in rare cell subsets (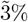 of total cells, Fig. 3) and effectively captures non-contiguous patterns across disjoint neighborhoods (Fig. 4B) by leveraging global manifold transport. Furthermore, significant shifts in cell type abundance naturally contribute to the differential signal. This ensures that the emergence or depletion of a cell state carrying a specific module is correctly flagged as a differential event.

There remain opportunities for further refinement. One important direction is to incorporate multiple biological replicates, which improves the statistical reliability of detected genes and modules, thereby enabling more rigorous cross-condition comparisons. In addition, while we have adopted computational strategies from GeneTrajectory to accelerate pairwise optimal transport computations, the method remains computationally demanding for large gene sets, warranting continued optimization. Finally, while our current work focused on scRNA-seq data, the concept of co-localization naturally extends to spatial transcriptomics. Measuring gene co-localization directly across spatial coordinates can reveal condition-specific spatial patterns of gene organization, providing a robust framework to link gene-level coordination with spatially organized biological processes.

## 4 Methods

### 4.1 DiCoLo Workflow

#### Input data

Let 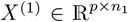 and 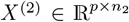 denote the log-normalized gene-by-cell expression matrices for the two conditions. An element 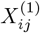 represents the normalized expression of gene *g*_*i*_ in cell *c*_*j*_ in condition 1. The two matrices are obtained by selecting a subset of informative genes for each condition, for example, based on their variability across cells. The union between the two gene subsets determines the gene set used as input to our method.

#### Gene–gene optimal transport (OT) distance

For each condition, DiCoLo computes a gene–gene distance matrix *D*^(1)^, *D*^(2)^ ∈ ℝ^*p×p*^ that measures dissimilarity between genes based on their expression distributions across cells. The optimal transport (OT) distance is computed following the procedure described in GeneTrajectory[8]. For completeness, we provide a brief description here. The computation of the OT distance is done in two steps:

- **Computation of the cell graph**. For each condition, we construct a k-nearest-neighbor (kNN) graph whose nodes correspond to individual cells. The k-neighborhoods are computed based on a low-dimensional representation of the cell geometry (e.g., diffusion maps). We denote by 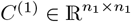, and 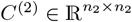 two cost matrices whose elements 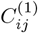 and 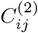 contain the shortest path distance from cell *i* to *j* in the condition 1 and 2, respectively.
- **Computation of OT distances between pairs of genes**. Let 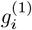 and 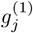 denote the gene–expression vectors of genes *i* and *j* over the cells of condition 1, normalized to sum to one. We denote by 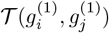 a group of matrices that satisfy the following constraints,

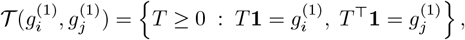

where **1** is the all-one vector. The group 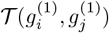 represents all potential transport plans from 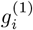 to 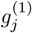. We compute the OT distance between these distributions by solving

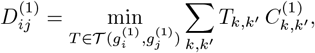

where *C*^(1)^ is the cell–cell cost matrix computed in the first step. A parallel formulation is used to obtain 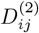 for condition 2.

The hyperparameters used to compute OT distances (e.g., the number of neighbors (*k*) used for the cell graph) follow the default settings in GeneTrajectory [8]. See Methods-’Robustness evaluation’ for a detailed evaluation of robustness to graph construction parameters.

#### Differential graph operator

##### Affinity and normalized graph construction

For each condition, the gene-gene OT distance matrix is converted into a symmetric affinity matrix *K* using a local-adaptive Gaussian kernel:

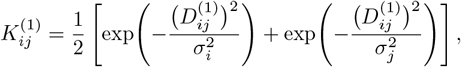

where *σ*_*i*_ is the distance from gene *i* to its *k*-th nearest neighbor. The choice of *k* follows the adaptive kernel bandwidth strategy described in [32]. We denote by *A*^(1)^ a diagonal matrix containing the degrees of the graph, such that 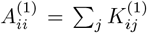. The kernel matrices are normalized to define the two graph operators *L*^(1)^ and *L*^(2)^ corresponding to the two conditions:

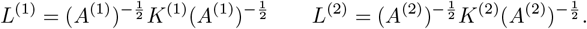

##### Rationale and differential operator formulation

To systematically detect changes in gene co-localization, we formulate the problem as identifying signals that are densely connected in the graph of one condition but sparsely connected in the other. The quadratic form *v*^*T*^ *L*^*k*^*v* acts as a score for the connectivity strength of a gene weight vector *v* on the graph of condition *k*. A high value of this quadratic form indicates that the genes assigned large weights in *v* are strongly co-localized within condition *k*. We therefore seek gene weight vectors *v* that reveal strong co-localization in condition 1 relative to condition 2. Mathematically, this is achieved by maximizing the generalized Rayleigh quotient:

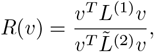

where 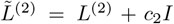. The regularization constant *c*_2_ serves as a spectral filter to prevent non-dominant components of *L*^(2)^ from being amplified during the optimization. It is set to the eigenvalue (*λ*_*ℓ*_) at the knee point (index *ℓ*) of the eigenvalue spectrum of *L*^(2)^. The specific method for determining the knee point is described in Supplementary Methods. Maximizing this ratio naturally selects vectors *v* for which connectivity in condition 1 is significantly stronger than in condition 2. The solution is given by the generalized eigenvalue problem 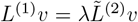. While this theoretically corresponds to the non-symmetric operator 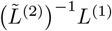, in practice, we utilize the symmetric differential operator which shares the same spectrum but offers superior numerical stability:

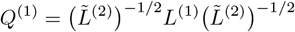

The eigenvectors of *Q*^(1)^ corresponding to the largest eigenvalues identify genes specifically co-localized in condition 1. Symmetrically, to capture genes co-localized in condition 2 relative to condition 1, we define the complementary operator 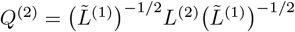.

#### Spectral analysis for differential co-localization gene extraction

The differential operators *Q*^(1)^, *Q*^(2)^ are subjected, separately, to eigenvalue decomposition:

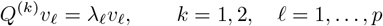

yielding eigenvectors *v*_*ℓ*_ that represent differential co-localization components and corresponding eigenvalues *λ*_*ℓ*_ that quantify their relative significance. Eigenvalues are sorted in descending order, and significant components are selected by identifying the knee point of the eigenvalue spectrum within the first 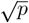 eigenvalues, where *p* is the total number of eigenvalues, to avoid numerical artifacts in the flat tail of the spectrum. The eigenvectors corresponding to the selected eigenvalues capture the dominant differential co-localization structure. For each selected eigenvector, loadings are standardized by z-transformation. Local false discovery rate analysis is performed using the R package `locfdr` (v.1.1.8)[33], assuming a standard normal null distribution (`nulltype = 0`). Genes with empirical FDR values below the expected FDR threshold returned by `locfdr` are defined as significant genes.

### 4.2 Downstream analysis: Gene module Assembly and Visualization

#### Gene modules assembly

Condition-specific gene modules are obtained by clustering the significant genes based on their differential operator on their respective OT distance submatrix. For example, significant genes from a vector of *Q*^(1)^ are clustered according to a submatrix of *D*^(1)^ that contains the OT between these genes. Hierarchical clustering with average linkage is performed using the `hclust` function (R package `stats`, v4.4.0[34]), and clusters are obtained with the dynamic tree-cutting algorithm (`cutreeDynamic` function from R package `dynamicTreeCut`, v1.63.1[35]). Genes with mean intra-module distances greater than median + 3 × MAD are excluded as outliers. Only modules containing at least 5 genes are retained for downstream analysis.

Module importance is calculated by weighting each module according to the spectral rank of its components. Each gene is uniquely assigned to its highest-ranked contributing eigenvector. For module *m*, we calculate *c*_*m,ℓ*_, the fraction of its genes assigned to eigenvector *ℓ*. The overall importance score is then computed as the weighted sum of these fractions: 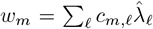, where 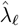 is the normalized eigenvalue. This metric prioritizes modules dominated by genes driven by the most significant spectral signals.

#### Module Activity Scores

Module activity scores are computed using `AddModuleScore` function (R ackage `Seurat`, v5.1.0[36]), then normalized across cells to have zero mean and unit variance, followed by min-max scaling to the [0, 1] range.

### 4.3 Experiments and analyses

#### Simulation workflow

We design a simulation framework that introduces synthetic co-localized expression patterns into real single-cell RNA-seq datasets. Each dataset contains two conditions or batches that either represent randomly split populations or real experimental batches with potential technical variation (see Methods-’Dataset and Processing’). One condition is used as the reference, while the other is modified by introducing artificial co-localization signals. Each introduced signal represents a gene group whose expression levels are modified to exhibit coordinated activation within a cell neighborhood, defined as a subset of transcriptomically similar cells. Multiple neighborhoods with varying sizes and gene groups are simulated for each dataset, each following the same procedure but with an independent random seed.

##### Definition of cell neighborhoods and gene selection

Each co-localization signal is generated by defining a cell neighborhood and selecting a subset of genes for perturbation. A neighborhood is created by randomly selecting a core cell and identifying its nearest neighbors based on Euclidean distance on the coordinates of the top 10 PCs. For each neighborhood, a subset of genes is randomly chosen for perturbation. Candidate genes satisfy two criteria: (1) they are expressed in 0.5–50% of cells and detected in at least one cell of the other condition, and (2) they show variance greater than mean expression, consistent with negative-binomial-like dispersion.

##### Generation of co-localized expression signals

We then simulate gene expression patterns that are confined to the cell neighborhood directly on the raw count data scale.

For each selected gene, the expression values of cells within the neighborhood are assigned values drawn from

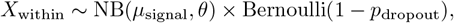

to mimic locally increased expression, while cells outside the neighborhood are assigned values drawn from

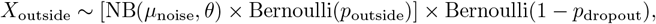

to introduce background noise. Here, the dispersion parameter (*θ*) is set to the median of gene-wise dispersions estimated from the raw count data, and the mean parameters, *µ*_signal_ and *µ*_noise_, are set to the 95th percentile and median of gene-wise means, respectively. The background expression fraction (*p*_outside_) specifies the fraction of cells outside the neighborhood allowed to express a perturbed gene. It is set to the smaller of two values: a baseline fraction (default 0.3) and the ratio of the neighborhood size to the total number of outside cells. This ensures that the number of expressing cells outside the neighborhood never exceeds that inside, thereby strictly enforcing the necessary localization of the signal. Finally, *p*_*dropout*_ represents the dropout probability.

Notably, expression values are sampled independently for each gene. This creates a challenging benchmark where co-localization is driven solely by spatial constraints without the aid of cell-to-cell covariance. Since explicit covariance would further increase the distributional similarity (reducing OT cost), this setup represents a conservative lower bound on the method’s sensitivity.

#### Dataset and Processing

Three real single-cell RNA-seq datasets (Human pbmc10k, Human pancreas, and Human pbmcsca) are used to perform simulations by perturbing the expression of selected genes while preserving realistic expression variability and incorporating realistic batch effects.

##### Human pbmc10k

The pbmc10k dataset is used[37]. CD14^+^ monocytes are extracted and randomly split into two subsets (1424 and 1423 cells) to create two pseudo-conditions without batch or biological differences.

##### Human pancreas

Two inDrops batches (“human2” and “human3”) from the human pancreas dataset [23] are used. Both batches are generated in the same laboratory using the same sequencing platform. Alpha cells are selected for analysis, and to minimize biological heterogeneity across subtypes, we follow [15] to retain the largest matched clusters from each batch (596 and 1023 cells). Genes expressed in fewer than 1% of cells are removed, and the larger batch is downsampled to 596 cells to maintain balanced sample sizes.

##### Human pbmcsca

Two batches (“Drop-seq” and “inDrops”) from the human pbmcsca dataset[24] are used. Both batches are derived from the same donor (PBMC1) within the same experiment, but are sequenced on different platforms. Cytotoxic T cells are selected for analysis, yielding 1189 and 1108 cells from the Drop-seq and inDrops batches, respectively. Genes expressed in fewer than 1% of cells are removed, and the larger batch is downsampled to 1108 cells to maintain balanced sample sizes.

##### Mouse embryo skin data

Publicly available mouse embryonic skin datasets, including E13.5 SmoM2 with paired wild-type (WT) controls[28] and E14.5 Wls with paired WT controls[8], are analyzed. Cell-type annotations for SmoM2 are stratified based on the expression of canonical dermal markers (*Sox2, Lef1, Dkk2*), while the original dataset provides annotations for Wls.

For all datasets, genes expressed in 0.5–50% of cells and ranked among the top 500 highly variable genes (HVGs) within each sample are retained. Genes detected in fewer than one cell in either sample are excluded. The union of the filtered gene sets from both samples is used for pairwise gene–gene OT distance computation, and in simulations, perturbed genes are additionally included. All datasets are processed using Seurat v5 [36]. Expression matrices are log-normalized, the top 2000 HVGs are selected, and the matrices are then scaled. Principal component analysis (PCA) is performed, and the top 10 PCs are used as input for the `GeneTrajectory` R package[8] to compute pairwise gene OT distances with default parameters. The resulting distance matrices are converted to affinity graphs using a Gaussian kernel (locally adaptive, k=10). For the SmoM2 and Wls datasets, gene modules are identified using `cutreeDynamic`[35] with `minClusterSize = 10` and `deepSplit = 0`.

#### Benchmarking with existing methods

We benchmark DiCoLo against representative methods for differential co-expression (DGCA) and cluster-free single-cell DE testing (miloDE and LEMUR) using the simulations described in Methods - ‘Simulation Workflow’ and ‘Dataset and Processing’. All methods are applied to the same gene set and run with default parameters unless otherwise specified. For miloDE and LEMUR, which require multiple replicates per condition, each condition is randomly split into two pseudo-replicates to enable comparison.

- **DiCoLo**: Genes are ordered by the absolute value of the first eigenvector loading of the differential operator *Q*^(1)^ or *Q*^(2)^, where *Q*^(1)^ or *Q*^(2)^ is constructed depending on the testing condition.
- **DGCA**: DGCA tested differences in pairwise gene–gene correlations between two conditions. Gene pairs that are positively correlated in the testing condition and show differential correlation in the same direction are selected. Each gene is ranked by the smallest adjusted *p*-value among its selected pairs; if multiple genes have the same adjusted *p*-value, they are ranked by descending order of the differential correlation statistic (`zScoreDiff`).
- **LEMUR**: LEMUR identifies, for each gene, a compact neighborhood showing consistent DE patterns. Genes upregulated in the testing condition are selected and ranked in ascending order of adjusted *p*-value; if multiple genes have the same adjusted *p*-value, they are further ordered by descending log fold change.
- **miloDE**: Following the authors’ recommendation, the two conditions are integrated using the “supervised” approach - Azimuth[38]. miloDE defines neighborhoods and performs DE tests within each neighborhood. Genes upregulated in the testing condition are selected and ranked in ascending order of their smallest adjusted *p*-value across all neighborhoods; if multiple genes have the same adjusted *p*-value, they are further ordered by descending log fold change.

#### Normalized AUPRC

Let AUPRC denote the raw area under the precision-recall curve and *p* denotes the positive fraction, *p* = *N*_*pos*_*/*(*N*_*pos*_ + *N*_*neg*_). The normalized AUPRC is defined as 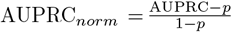. This rescales the metric to range between 0 and 1, where 0 corresponds to random performance, and 1 indicates perfect detection.

#### Robustness evaluation

DiCoLo relies on a cell graph for each condition, constructed from *k*-nearest neighbors in a dimensionality-reduced embedding. By default, we follow the parameter selection strategy from GeneTrajectory: we select the number of non-trivial diffusion map components by identifying where the eigenvalue spectrum begins to flatten (typically 10-15 components) and set *k* = 10. To evaluate the sensitivity of DiCoLo to graph construction parameters, we systematically vary three components using the SmoM2 dataset: the number of nearest neighbors (*k*), the manifold dimensionality, and the embedding method. First, we test manifold dimensionality by varying the number of diffusion map components from 5 to 25 while fixing *k* = 10. Second, we vary *k* from 5 to 25 while holding the cell embedding fixed at 10 diffusion map components. Third, we compare the default diffusion map against an alternative PCA-based embedding using the top 10 PCs (with *k* = 10). For each setting, we evaluate the stability of identified differentially co-localized genes by computing the Jaccard index between gene sets derived from the leading eigenvectors of the differential operator (Fig. S6).

#### Computational time

We evaluate the runtime of DiCoLo using the mouse SmoM2 dataset. After selecting the top 500 HVGs in each condition, we obtained a merged set of 712 genes. The analysis proceeds in two stages: (i) computation of pairwise gene–gene OT distances and (ii) construction of differential graph operators followed by spectral analysis. The computational cost is dominated by the first stage. We assess performance on a Linux system equipped with AMD EPYC 7643 processors, restricting the analysis to 8 parallel processes. For the condition containing 8,975 cells, the OT computation required 671.3 seconds, while the condition with 3,663 cells required 316.2 seconds. In contrast, the second stage was completed in approximately 2.1 seconds. For significantly larger datasets (e.g., *>* 20, 000 cells or *>* 2, 000 genes), we recommend applying the cell graph coarse-graining and gene affinity graph sparsification strategies utilized in [8].

### 4.4 Data availability

The human PBMC10k dataset is available at https://support.10xgenomics.com/single-cell-gene-expression/ datasets/3.0.0/pbmc_10k_v3. The human pancreas and PBMCSCA datasets are included in the SeuratData package and can be accessed as panc8.SeuratData and pbmcsca.SeuratData from https://github.com/satijalab/seurat-data. The mouse embryonic skin datasets are available under GEO accessions GSE280825 (E13.5 SmoM2) and GSE255534 (E14.5 Wls).

### 4.5 Code availability

The R package of DiCoLo and the code used for data analysis are available on GitHub (https://github.com/KlugerLab/DiCoLo).

## 4.6 Acknowledgment

The authors thank Ronald Coifman, Peggy Myung, Boris Landa, Fabio Parisi, Francesco Strino, Ri-hao Qu, and members of the Kluger Lab for fruitful discussions. Y.K. is supported by [R01GM131642, R01DA063148, UM1PA051410, U54AG076043, U54AG079759, U01DA053628, P50CA121974, R33DA047037].

## Notes

### Competing Interest Statement

The authors have declared no competing interest.

### Summary of Updates

Methods-Differential graph operator Methods-Robustness evaluation Methods-Computational time

