## Supplementary Material for "DiCoLo: Integration-free and cluster-free detection of localized differential gene co-expression in single-cell data"

### Supplementary: DiCoLo: Differential co-localization analysis

#### S1. Supplementary Figures

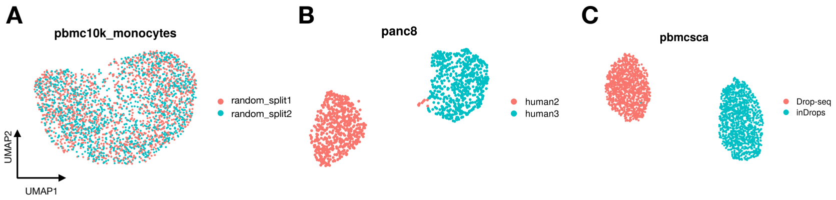


**Fig. S1.** UMAP of each dataset with two batches, generated using the top 2,000 highly variable genes. (A) human PBMC10k, (B) human pancreas, and (C) human PBMCSCA, color-coded by batch.

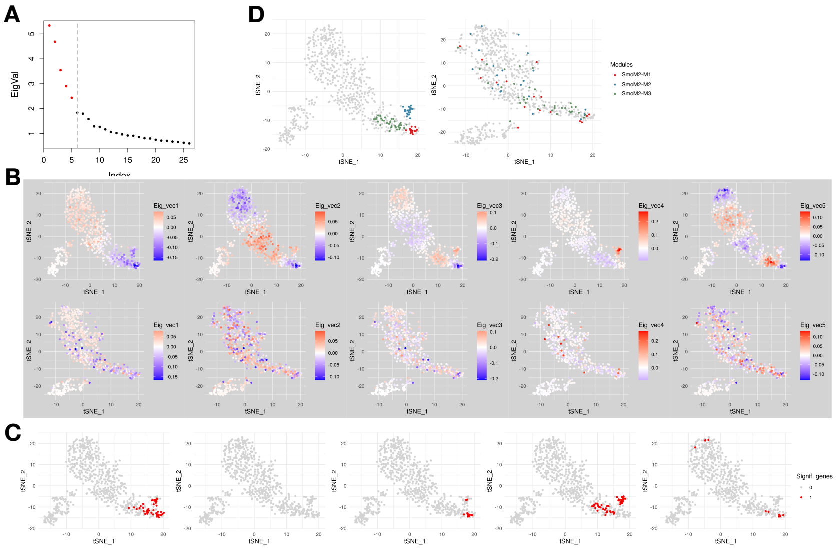


**Fig. S2.** Derivation of co-localized gene modules of SmoM2 (compared with CTL). (A) Eigenvalue spectrum of the differential graph operator for SmoM2, with red points marking the top eigenvalues selected at the knee point. (B) Gene embeddings generated by t-SNE based on OT distances (first row SmoM2, second row CTL), color-coded by eigenvector loadings. Each column corresponds to one of the top eigenvectors selected in (A). (C) Gene embeddings of SmoM2, with significant genes (in red) determined by the locFDR threshold, as detailed in Methods - 'Gene modules assembly'. (D) SmoM2-specific co-localized gene modules obtained by hierarchical clustering of significant genes in (C) based on SmoM2 OT distances, overlaid on SmoM2 (left) and CTL (right) gene embeddings. Parameter details are provided in Methods - 'Gene modules assembly' and 'Mouse embryonic skin data'

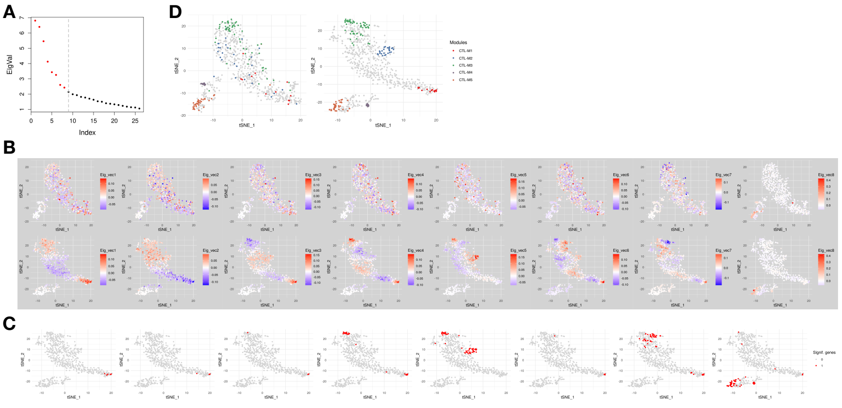


**Fig. S3.** Derivation of CTL-specific co-localized gene modules (compared with SmoM2). (A) Eigenvalue spectrum of the differential graph operator for CTL, with top eigenvalues in red selected at the knee point. (B) Gene embeddings of SmoM2 (first row) and CTL (second row), color-coded by loadings of the top eigenvectors selected in (A). (C) CTL gene embedding with significant genes highlighted in red by the locFDR threshold. (D) CTL-specific co-localized gene modules obtained by hierarchical clustering of significant genes in (C), overlaid on SmoM2 (left) and CTL (right) embeddings.

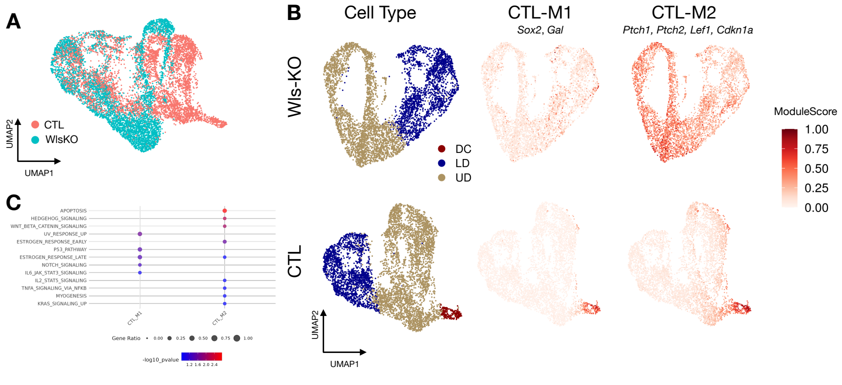


**Fig. S4.** DiCoLo reveals condition-specific co-localized gene modules in DC genesis (Wls vs wild-type control). (A) UMAP of cells color-coded by condition. (B) UMAP of cells embedded separately for each condition, color-coded by cell type (left) and by co-localized gene modules (right), including two control-specific modules. (C) MsigDB pathway enrichment analysis of these gene modules, showing the top 5 enriched pathways ($p$ value $< 0.1$) for each module.

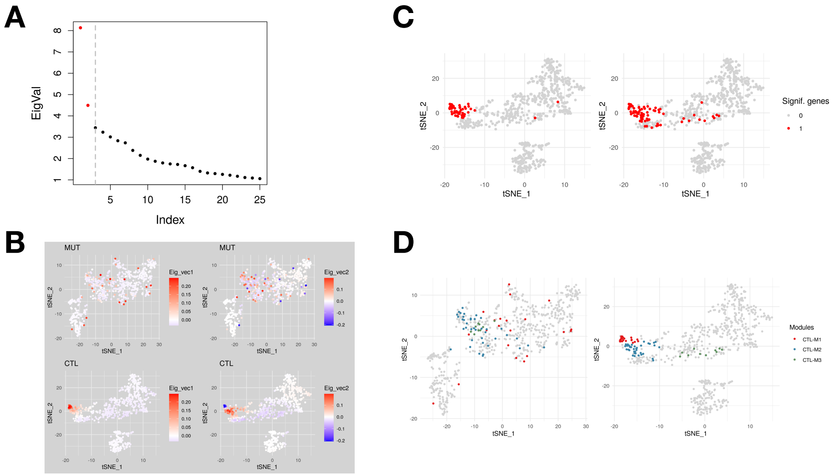


**Fig. S5.** Derivation of CTL-specific co-localized gene modules (compared with Wls). (A) Eigenvalue spectrum of the differential graph operator for CTL, with top eigenvalues in red selected at the knee point. (B) Gene embeddings of Wls (first row) and CTL (second row), color-coded by loadings of the top eigenvectors selected in (A). (C) CTL gene embedding with significant genes highlighted in red by the locFDR threshold. (D) CTL-specific co-localized gene modules obtained by hierarchical clustering of significant genes in (C), overlaid on Wls (left) and CTL (right) embeddings.

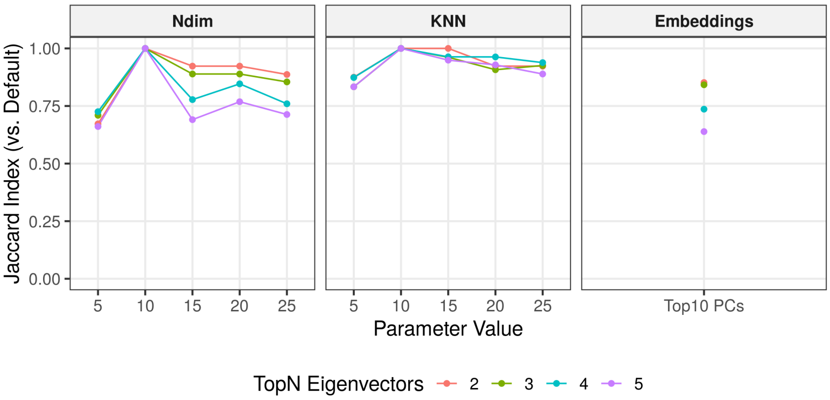


**Fig. S6.** Robustness of DiCoLo to graph construction parameters. (Left, Middle) Stability of identified differentially co-localized genes measured by the Jaccard index between candidate genes identified using the default cell graph settings (top 10 diffusion map components, $k = 10$) and those using varying number of diffusion map components (Left) and neighborhood size (Middle). (Right) Jaccard index comparing the default top 10 diffusion map components against top 10 principal components. Colors denote the number of leading eigenvectors of the differential operator used for gene selection. Analyses were conducted on the SmoM2 dataset and test the differential operator of SmoM2 against CTL.

#### S2. Supplementary Methods

**Knee-point selection of informative eigenvectors** Eigenvalues are sorted in descending order, treated as a one-dimensional curve, and a straight line is drawn between the first and last points. For each eigenvalue, we compute its perpendicular distance to this line; the index with the largest distance is selected as the knee point, marking the transition from signal-dominated to noise-dominated components.

**Pathway enrichment analysis** Pathway enrichment is performed using the `msigdbr` R package (v7.5.1)[1] and `clusterProfiler` R package (v4.14.6)[2], based on the Hallmark gene sets from the Molecular Signatures Database (MSigDB)[3]. For each gene module, the top five enriched pathways with $p$-values $< 0.1$ are reported.

#### S3. Supplementary Data

Supplementary Data can be downloaded from: `https://github.com/ruiqi0130/DiCoLo_supplementary_data/raw/main/Supplementary_data.xlsx`. which includes:

- Supplementary Data1: scRNA-seq datasets used in this paper.
- Supplementary Data2: Co-localized gene modules identified from mouse hair follicle datasets.
